# Near chromosome-level genome assembly for the invasive annual forb *Centaurea melitensis*

**DOI:** 10.64898/2026.05.18.726060

**Authors:** Anthony J. Dant, Jessie A. Pelosi, Poppy C. Northing, Katrina M. Dlugosch

## Abstract

**Premise:** *Centaurea melitensis* (Asteraceae) is a problematic invader of grasslands globally, but little is known about its genetic makeup. Here we develop a reference genome to facilitate studies of its invasion history, genetic variation, and evolution.

**Methods:** Inbred offspring of a single individual of *C. melitensis* from its invasion of California, USA were used for flow cytometry to estimate genome size, and for genomic DNA extraction. DNA was sequenced with PacBio HiFi technology (yield = 85.7Gb). The genome was assembled with Hifiasm and annotated with BRAKER3. GENESPACE was used to compare gene order (synteny) with three other species within the subfamily Cichorioideae.

**Results:** We estimated a mean genome size of 795.0 Mbp for *C. melitensis*, and our assembly totaled 696.6 Mbp in 48 contigs (N50 = 55.6 Mbp; BUSCO = 98%), with annotation of 25,157 protein-encoding genes. This included four telomere-to-telomere putative chromosomes, nine additional chromosome arms terminated by telomeric repeats, and a complete chloroplast genome. Synteny varied markedly across the genus and subfamily, suggesting a dynamic history of structural variation in the lineage of *C. melitensis*.

**Discussion:** We provide a highly complete and contiguous genome assembly to facilitate the further study of genomic variation in *C. melitensis*.

## INTRODUCTION

*Centaurea melitensis* L. (Asteraceae), commonly known as Maltese starthistle or tocalote, is an annual forb that is native to the Mediterranean region across Southern Europe and Northern Africa (DiTomaso and Healy 2007). This plant is also a prolific invasive species that is now found on every continent in the world except Antarctica (Hellwig 2003). *Centaurea melitensis* currently exists in its highest densities in its invaded ranges in South America, primarily in Chile (Sotes et al. 2021), and in North America, primarily in California, USA (Moroney and Rundel 2013a). Populations in North America appeared with Spanish colonization, with the oldest specimen of *C. melitensis* found within an adobe brick dated to 1797 in San Fernando, California (Hendry 1931). It is classified as both an agricultural pest for its degradation of pasture land and an ecological danger for its ability to outcompete and displace native annuals (Moroney and Rundel 2013b; Roché and Roché 2021; California Invasive Plant Council 2026).

Several potential reasons for *C. melitensis*‘s invasion success have been hypothesized, most related to its ability to successfully grow in a variety of environments. The species is primarily self-fertilizing, with both chasmogamous and cleistogamous capitula and it has achene heteromorphism (Porras and Muñoz 2000a), suggesting an ability to reproduce without needed interactions from other organisms when colonizing novel environments. It has been found to occupy a wide variety of different habitats including chaparral, oak woodland, grassland, montane, and urban environments (Roché and Roché 2021), and it will readily colonize post burned areas (Riba et al. 2002; Moroney and Rundel 2013b). Additionally, *C. melitensis* shows both local adaptation and plasticity in response to variation in soil nitrogen deposition (Moroney et al. 2013) and aridity (Humphries et al. 2024), again suggesting ecological flexibility, though this may vary across its invaded range (Bozzolo and Lipson 2013; Valliere et al. 2025). The species has also been linked to specific aspects of urbanization such as herbicide usage and impervious surface in California, suggesting the potential for urban environments to be centers of dispersal for the species (Dant et al. 2025).

Despite a history of research interest in the adaptability of *C. melitensis* and its invasion success across the globe, little is known about its genetics or genomics. The species has been included in phylogenetic surveys of the genus *Centaurea*, with Garcia-Jacas et al. (2006) most recently placing it in a western Mediterranean ‘Hymenocentron-Melanoloma-Seridia group’ of congeners, with *C. melitensis* most closely related to another well-known invasive species *C. solstitialis* L. (yellow starthistle), among the species included. The base chromosome number for this clade is 1N = 11 (Garcia-Jacas et al. 2006), but *C. melitensis* has been reported as 1N = 9, 12, and 18 (Keil 2012) and *C. solstitialis* as 1N = 8 (Widmer et al. 2007), indicating karyotypic evolution and variability within both the clade and species. We are unaware of any other genetic studies of *C. melitensis*. Genomic analyses in particular would be powerful for identifying routes of invasion and the genetic basis, variability, and evolution of traits underpinning invasion success (Welles and Dlugosch 2018; Sherpa and Després 2021; McGaughran et al. 2024; Hodgins et al. 2025). For example, genetic and genomic analyses in closely-related C. *solstitialis* have uncovered the routes and timing of its introductions and trait evolution during invasion (Barker et al. 2017; Irimia et al. 2023), the genetic basis of invasion-associated traits (Reatini et al. 2024), and evidence of genome size evolution during invasion (Cang et al. 2024).

Here we provide the first assembled and annotated genome of *C. melitensis* to facilitate the genomic study of this invader. Using tissue of inbred offspring from the California invasion, we used flow cytometry to provide the first estimate of a genome size for *C. melitensis*. Then, we sequenced high molecular weight total DNA using a PacBio HiFi long read library on a single Revio SMRT flow cell. With these data, we assembled highly complete nuclear and chloroplast reference genomes from these reads, and annotated their contents. We report genome content and evidence of dynamic evolution of gene order (synteny) and karyotype between *C. melitensis* and its near and distant relatives across the Asteraceae. This resource can be used to better understand the evolution of the *C. melitensis* genome and the genetic mechanisms responsible for its invasion around the world.

## METHODS

### Reference tissue

We obtained leaf and root tissue for flow cytometry and genomic sequencing from progeny of an individual of *C. melitensis* collected near Sandstone Peak in the Santa Monica Mountains (California, USA) in summer 2021 (34°06’26“N 118°54’33“W) (Figure 1). Progeny of this individual were allowed to self-fertilize in a greenhouse for two generations. We placed multiple third generation seeds of full siblings on soil in petri dishes indoors under 12 h of fluorescent lights at 21°C until they germinated. After one week of growth, we transferred seedlings to commercial potting soil (Happy Frog Potting Soil, FoxFarm Soil and Fertilizer Co, Arcata, CA, USA) in D25L deepots (Steuwe and Sons, Tangent, Oregon, USA) in a greenhouse at the University of Arizona (Tucson, Arizona, USA) where they were hand watered under ambient light. Plants were grown in the greenhouse for three months from November to January. Leaf and root tissue for DNA sequencing were collected from a single individual, immediately flash frozen in liquid nitrogen, and stored at −80 ℃ until DNA extraction (see ‘*DNA Extraction and Sequencing*’ below). A sibling of this individual is accessioned at the University of Arizona Herbarium (ARIZ; accession # pending).

**Figure 1.**
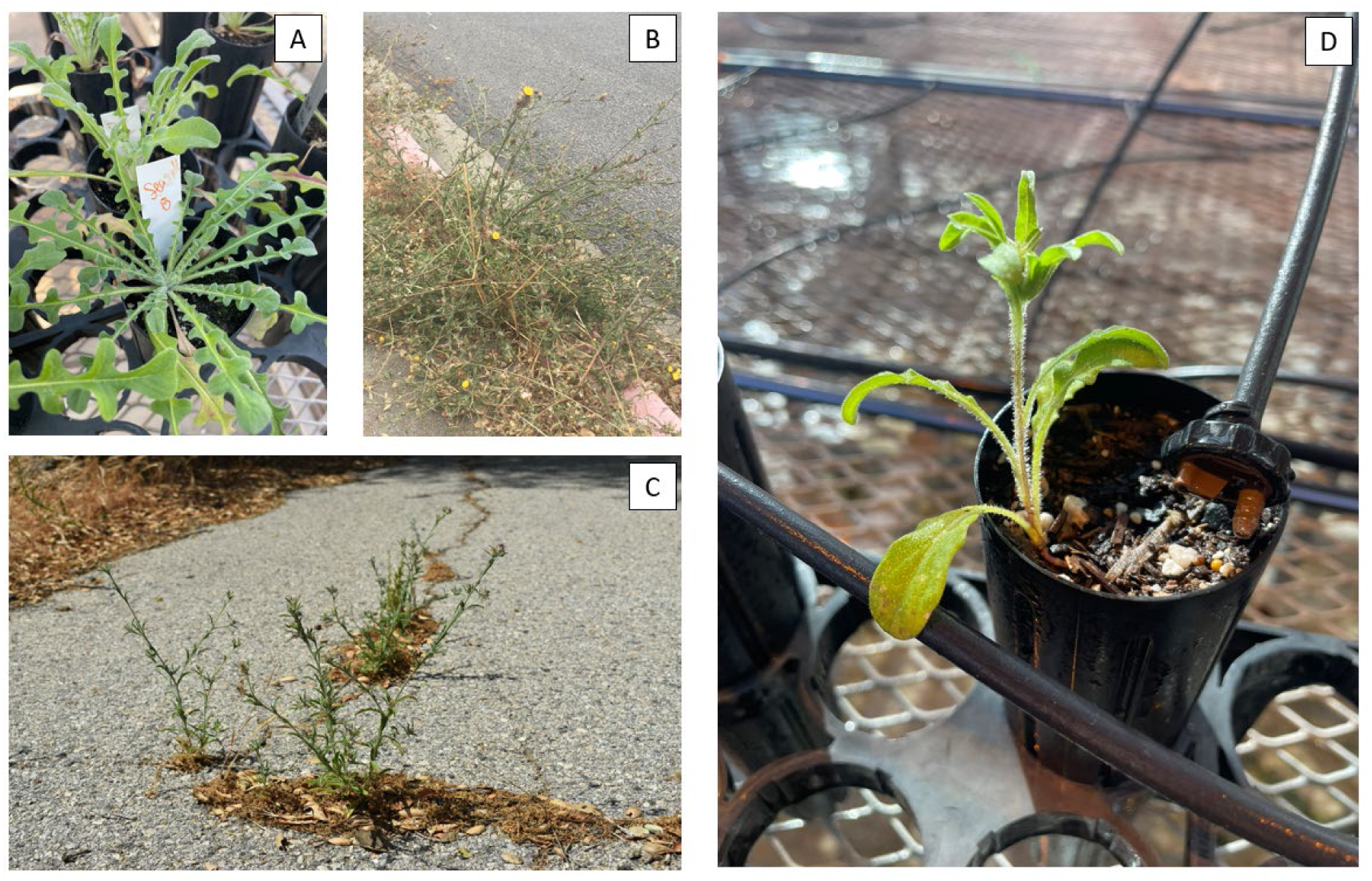
Images of *Centaurea melitensis* A) producing rosette leaves while growing at the University of Arizona greenhouse, B) flowering while growing out of a sidewalk in Monterey, California, and C) growing in a concrete crack near Malibu Creek State Park, California. D). The individual that was sequenced for this assembly.

### Genome Size Estimation

We used flow cytometry to estimate the genome size of a three-month-old full sibling of the individual sequenced for the genome assembly, following the two-step protocol developed by (Dolezel, Greilhuber, and Suda 2007) with modifications described in (Cang et al. 2024; Northing et al. 2025). We measured the 2C genome content of five different leaves from this individual on two separate days using a FACSCanto II flow cytometer (BD Biosciences, San Jose, California, USA). In brief, nuclei were isolated from each leaf (mass ≈ 60 mg) by co-chopping with 50 mg of leaf tissue from *Raphanus sativus* ‘saxa’ (2C = 1.11pg, provided by the Institute of Experimental Botany, Prague, Czech Republic), which served as an internal standard. Leaves were co-chopped in 500 µL of Otto I buffer on an ice-cold glass petri dish, using a fresh razor blade for each leaf. The nuclear suspension was filtered through 18µm Nylon mesh into a 5 mL polystyrene round-bottom flow cytometry tube and kept on ice. We also prepared a sample with only *R. sativus* each day of analysis to validate the identification of the standard and sample peaks. Once all samples were prepared, we added 770 µL of Otto II buffer and 7.5 µL of 1mg/mL RNAse to each suspension, which we then incubated for 5 minutes. After incubation, we added 29 µL of 1mg/mL propidium iodide (PI) staining solution to each sample. We covered the samples to shield them from light and allowed them to incubate for another 5 minutes before analyzing on the FACSCanto II with a low flow rate. We calculated the 1C genome size using the G1 peaks of mean PI fluorescence from the sample and standard following the formula described by Dolezel and Bartos (2005). Every peak used to calculate a genome size had a coefficient of variation less than 5% (Table S1 in Appendix S1). We then compared this genome size estimate to those published for ten other *Centaurea* species (see Table S2 in Appendix S1).

### DNA Extraction and Sequencing

DNA extraction and sequencing were performed by the Arizona Genomics Institute (AGI; Tucson, Arizona, USA). High molecular weight DNA was extracted with the Macherey-Nagel NucleoBond kit (Macherey-Nagel, Allentown, PA, USA) from 0.4 g of flash-frozen tissue ground in liquid nitrogen with a mortar and pestle. The eluted DNA was then precipitated with cold isopropanol. DNA was collected by centrifugation at 4500 g for 20 minutes, washed with 70% ethanol, and air dried. The pellet was resuspended thoroughly in 50 µl 10 mM Tris-HCl, pH 8. DNA purity was measured with a Nanodrop (Thermo Fisher Scientific, Waltham, MA, USA), concentration was measured with Qubit High Sensitivity kit (Invitrogen, Carlsbad, CA, USA), and fragment size was assessed by Femto Pulse System (Agilent, Santa Clara, CA, USA).

To prepare DNA for PacBio sequencing, DNA was sheared to 10-20 kb using a Megaruptor 3 (Diagenode, Denville, NJ, USA) followed by SMRTbell cleanup beads (Pacific Biosciences, Menlo Park, CA, USA). The sequencing library was constructed following manufacturers’ protocols using the HiFi prep kit 96 (Pacific Biosciences). The final library was size selected using AMPure PB beads (Pacific Biosciences). The recovered final library was quantified with the Qubit HS kit (Invitrogen) and size checked on the Femto Pulse System (Agilent). The final library was prepared for sequencing with the PacBio Revio SPRQ Sequencing Plate for HiFi libraries, loaded on a single Revio SMRT cell, and sequenced in CCS mode for 30 hours.

### K-mer Analysis and Genome Assembly

Before assembling the *C. melitensis* reference genome, we used *k*-mer analysis to estimate the genome size and level of homozygosity from the raw HiFi reads. We used KMC v3.2.4 (Kokot et al. 2017) to count *k*-mers for *k* = 17, 21, 25, 31, 41, 51, 61, 71, 81, 91, 101, 111, and 121 (with a maximum count threshold of 10,000 to exclude highly repetitive *k*-mers), and then used GenomeScope2 (Ranallo-Benavidez et al. 2020) to predict the genome size and homozygosity from the *k*-mer spectra.

We generated an initial de novo genome assembly with the raw HiFi reads using Hifiasm v0.25.0-r726 (Cheng et al. 2021). Due to low levels of measured heterozygosity found in the previous *k*-mer analysis and the known selfing mating system of the species (Porras and Muñoz 2000b), the -l0 flag was used to disable haplotype purging. We also generated a plastome assembly to identify and remove contigs from our initial genome assembly that likely represent plastid genome sequences. We mapped our raw HiFi reads to the *Carthamus tinctorius* L. reference chloroplast genome assembly (Lu et al. 2016) (GenBank accession: NC_030783.1) using Minimap2 v2.28 (Li 2018) with default settings. We then subsampled the mapped reads to 150x coverage (1,464 reads) using SeqKit v2.8.1 (Shen et al. 2016) with two-pass sampling and used those reads to assemble the *C. melitensis* plastid genome using Hifiasm v0.25.0-r726 (Cheng et al. 2021); this time with default settings. We then used Bandage v0.8.1 (Wick et al. 2015) to visualize the resulting assembly, which informed our decision to treat the longest contig in the assembly as the final *C. melitensis* plastome assembly (see Results). We then used GeSeq v2.03 (Tillich et al. 2017) with default parameters to annotate and visualize the plastid genome assembly and annotation.

To identify and remove chloroplast contigs from the nuclear assembly, we first separated contigs by size at a 1 Mbp threshold using SeqKit v2.8.1 (Shen et al. 2016); this threshold was based on chloroplast genome sizes in Asteraceae, which generally fall between 115,000 bps to 165,000 bps (Daniell et al. 2016). We then mapped small contigs (<1 Mb) to the *C. melitensis* plastome using Minimap2 v2.28 (Li 2018) with the asm5 preset. Contigs that mapped to the chloroplast were identified using SAMtools v1.10 (Li et al. 2009) and removed from the nuclear assembly. All remaining contigs that did not map to our plastome assembly were retained for the final assembly.

We used the purge_haplotigs pipeline v1.1.3 (Roach et al. 2018) to remove contigs representing haplotypic variation. HiFi reads were mapped to the assembly using Minimap2 v2.28 with the map-hifi preset, and contigs were classified using coverage thresholds of 10× (low), 80× (mid), and 160× (high). We also removed several other potential contaminants from the initial contigs using BlobToolKit v4.4.5 (Challis et al. 2020) with the bestsumorder taxonomic assignment rule to identify and remove contamination from other taxa in the genome assembly, and Inspector v1.31 (Chen et al. 2021) with the hifi datatype parameter to identify and correct any errors in the assembly. Thus, our final nuclear genome assembly for *C. melitensis* included all of the contigs that remained after filtering out putative organellar genome sequences, haplotypic variation, and contamination. We used BUSCO v5.8.3 (Tegenfeldt et al. 2025) with the eudicots_odb10 dataset (2020-09-10) to assess the completeness of our initial assembly and final assembly. Genome-wide heterozygosity was calculated by mapping HiFi reads to the assembled *C. melitensis* genome using Minimap2 v2.28 (Li 2018), calling variants with bcftools v1.19 (Danecek et al. 2021), and dividing the number of heterozygous SNPs by the total assembly length.

### Genome Annotation

To annotate repetitive elements in the assembly, we generated a custom repeat library for *C. melitensis* using RepeatModeler v2.0.3 (Flynn et al. 2020) with the -LTRStruct parameter, which utilizes LTR_retriever v3.0.2 (Ou and Jiang 2018) for structural identification of long terminal repeat (LTR) retrotransposons. Then we used RepeatMasker v4.1.8 (Smit et al. 2015) with soft-masking (-xsmall) to identify and mask repetitive elements in the final assembly using our species-specific repeat library as input. To identify telomeric repeats in the assembly, we ran tidk v0.2.0 (Brown, Manuel Gonzalez de La Rosa, and Blaxter 2025) with the --clade Asterales parameter to search for the telomeric repeat sequence (TTTAGGG)*_n_*. We used tidk ‘explore’ to identify potential telomeric regions with repeat lengths between 5 and 12 bp, followed by tidk ‘find’ to locate telomeric sequences.

To generate a structural gene annotation for *C. melitensis*, we used the BRAKER3 v3.0.6 pipeline (Gabriel et al. 2024) with multiple sources of extrinsic evidence, including RNAseq data from a close relative, *C. solstitialis,* and proteomes from nine species in Asteraceae (Table S3 in Appendix S1). We downloaded RNAseq data from six RNA libraries generated from *C. solstitialis* leaf tissue (Reatini et al. 2024; NCBI accession numbers (SRX18786794 - SRX18786799) and mapped the raw RNAseq reads to our soft-masked final genome assembly with hisat2 v2.2.1 (Kim et al. 2019) with default settings. Predicted proteomes for nine Asteraceae species were downloaded and provided to BRAKER3 for additional evidence (Table S3 Appendix S1). Using these data, we predicted gene models using the default BRAKER3 v3.0.6 pipeline (Gabriel et al. 2024), which relies on GeneMark-ETP (Bruna et al. 2024), DIAMOND (Buchfink et al. 2015), spaln2 (Gotoh 2008; Iwata and Gotoh 2012), StringTie2 (Kovaka et al. 2019), GFF utilities (Pertea and Pertea 2020), AUGUSTUS (Stanke et al. 2006, 2008), and TSEBRA (Gabriel et al. 2021). We used BUSCO v5.8.3 (Tegenfeldt et al. 2025) to assess the completeness of our gene annotation against the eudicots_odb10 dataset and OMArk v0.3.1 (Nevers et al. 2025) to assess annotation quality.

### Synteny analysis

We used GENESPACE v1.3.1 (Lovell et al. 2022) to examine macrosynteny among the genomes of four species in Asteraceae: *C. melitensis*, *C. solstitialis* (NCBI accession: GCA_030169165.1; Reatini et al. 2024), *Cynara cardunculus var. scolymus* L. (NCBI accession: GCF_001531365.2; Acquadro et al. 2020), and *Lactuca sativa* L. (NCBI accession: GCF_002870075.4; Reyes-Chin-Wo et al. 2017). GENESPACE uses OrthoFinder v2.5.4 (Emms and Kelly 2015, 2019) to cluster genes into orthogroups, identifies syntenic blocks with MCScanX v1.0.0 (Wang et al., 2012), and integrates these data to produce comprehensive visualizations of syntenic patterns across genomes. We included all contigs containing 100 or more annotated genes in the synteny analysis.

## RESULTS

### Flow cytometry and k-mer analysis

From flow cytometry, we estimated the 1C genome size for a full sibling of the individual used for our *C. melitensis* genome assembly as 0.812 ± 0.005 pg (795.04 ± 4.62 Mbp; Table S1 in Appendix S1). The PacBio Revio SMRT cell yielded a total of 85.7 Gb from 6,061,768 reads with a read N50 of 15.9 kbp. Our *k*-mer analysis of the raw PacBio HiFi reads yielded similar genome size estimates ranging from 563.81 Mbp (*k*=17) to 685.17 Mbp (*k*=121) across all *k*-mer sizes tested. A complete list of all *k*-mer outputs can be found within the Supplementary materials (Table S4 in Appendix S1; Figure S1 in Appendix S2). The median of published genome size estimates for ten other *Centaurea* species was 811.0 Mbp, with a range of 695.8 Mbp to 999.6 (Table S2 in Appendix S1).

### Chloroplast genome assembly

A total of 523,865 reads that mapped to the *Carthamus tinctorius* reference plastome were incorporated into the initial assembly of the chloroplast genome, with a mean length of 15,699 bp, totalling 8.224 Gb. Subsampling this to 1,464 reads (∼250x coverage), five contigs were assembled, which were visualized in Bandage v0.8.1. The largest contig was retained which totaled 161,562 bp, serving as our final plastome assembly. This plastome assembly had a typical quadripartite structure consisting of a large single-copy (LSC) region of 83,837 bp, a small single-copy (SSC) region of 18,586 bp, and two inverted repeats of 25,188 bp (IRa) and 33,951 bp (IRb), with a GC content of 37.98%. The annotation resulted in 121 genes, of which 83 were protein-coding genes, four were rRNAs, and 23 were tRNAs (Table S5 in Appendix S1; Figure S2 in Appendix S2).

### Nuclear assembly

The initial Hifiasm assembly contained 839 contigs totaling 734 Mbp with a contig N50 of 54.3 Mbp. BUSCO results from this initial assembly identified 2,279 (98%) complete orthologs from eudicots_odb10 with 2,150 (92.5%) being single-copy, and 129 (5.5%) duplicated. In addition, 21 (0.9%) were fragmented and 26 (1.1%) were missing (Table S6, Appendix S1). Following the removal of contigs that mapped to our assembled *C. melitensis* chloroplast genome, we retained all other contigs. This resulted in a total of 309 contigs. After removal of putative haplotypic variation, 48 contigs were retained as the final assembly. Blobtools identified no contamination in the final assembly (Figure S3, Appendix S2). Evaluating the final assembly with Inspector v1.31 (Chen et al. 2021) obtained a quality value score (QV) value of 64.31, indicating high assembly quality.

The final genome assembly with 48 contigs was 696,561,998 bp long, had a contig N50 of 55,586,964 bp and N90 of 37,081,860 bp. BUSCO identified 2,279 (98.0%) eudicot complete BUSCOs with 2,152 (92.5%) being single-copy, 127 (5.5%) duplicated, 21 (0.9%), fragmented and 26 (1.1%) missing (Figure 2, Table S6 in Appendix S1). The genome-wide heterozygosity was estimated to be 11.53 heterozygous SNPs per Mb.

**Figure 2.**
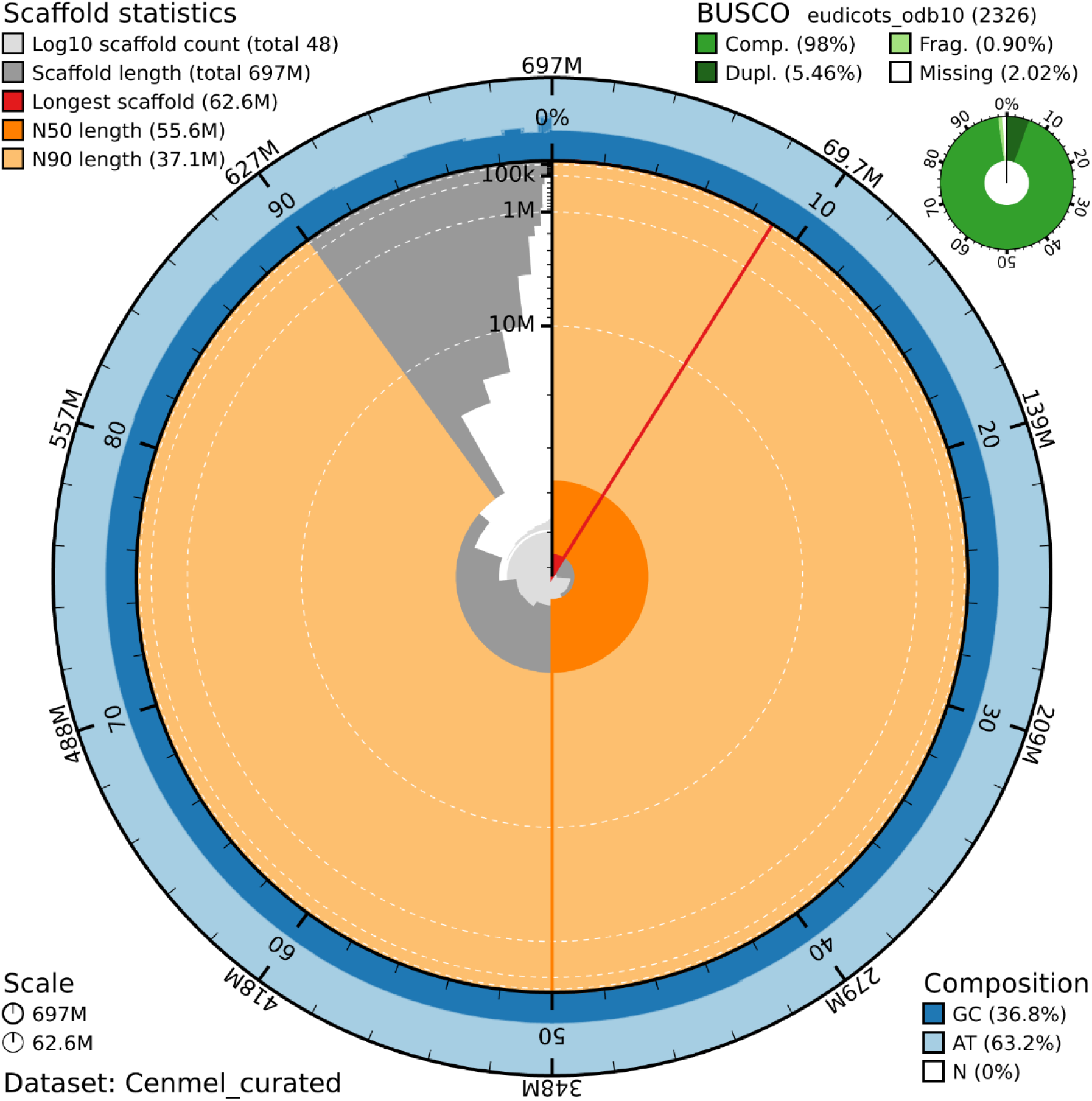
Blobtools snail plot of the final genome assembly statistics for *Centaurea melitensis*. The assembly consists of 48 contigs totaling 696.6 Mb. Colors in the graph represent the following: Dark and light blue represent GC (36.8%) and AT (63.2%) composition in the genome; the red line visualizes the length of the longest contig (62.6 Mb); light gray shows the log-scaled cumulative contig count while dark gray shows the distribution of total contig length; light and dark orange represent N50 (55.6 Mb) and N90 (37.1 Mb) lengths respectively; BUSCO results for the assembly are shown with complete BUSCOs in bright green (98%), duplicated BUSCOs in dark green (5.46%), fragmented BUSCOs in light green (0.90%), and missing BUSCOs in white (2.02%).

### Annotation

The final *C. melitensis* nuclear genome contained 65.85% repetitive content as identified by RepeatMasker. Of this, 64.19% was classified as interspersed repeats and 1.10% as tandem repeats, with an additional 0.37% corresponding to small RNAs. The interspersed repeats were dominated by long terminal repeats (LTRs; 26.61%) that mostly consisted of Ty1/Copia (11.52%) and Gypsy/DIRS1 (11.08%) elements. This was followed by unclassified elements (31.74%), DNA transposons (5.01%), and small proportions of LINEs (0.84%) and SINEs (0.003%) (Table S7 in Appendix S1).

Structural gene annotation using BRAKER3 yielded 25,157 predicted gene models encoding 27,382 transcripts, with a mean gene length of 3,767 bp, and an average of 5.4 exons per gene. Approximately 72.58% of these genes were multi-exonic. These genes collectively encode for 35.4 Mbp of coding sequence. We identified 93.1% complete eudicot BUSCOs in these genes (of which 83.1% were single-copy and 10.0% were duplicated) as well as 0.5% fragmented genes.. Functional domains were identified by InterProScan v5.66-98.0 (Joens et al., 2014) in 22,279 protein sequences (84.84% of the total structural gene models). OMArk v0.3.1 assessed annotation completeness at 93.57 %.

The largest 13 contigs contained the vast majority of genes, ranging from Contig 1 with 2,672 genes to Contig 13 with 1,007 genes; Table S8 in Appendix S1). The largest 17 contigs contained at least 100 genes each (the lowest being Contig 17 with 142 genes), suitable for synteny analyses (see below). Three other contigs contained less than 100 genes (range 1-69), and the remaining 28 contigs contained no genes in the annotation.

Using tidk v0.2.0, telomeric repeat motifs corresponding to the canonical Asteraceae telomere sequence (TTTAGGG)*_n_* ; Peska and Garcia 2020) were identified on 13 out of 17 of the largest contigs of our final genome assembly (Figure 3; Table S9 in Appendix S1). The repeat and its reverse complement were the most abundant tandem repeat motifs detected, occurring greater than 7,000 times across the assembly. Distinct telomeric blocks, each exceeding 400–900 tandem repeats, were detected at both ends of four contigs. The remaining nine contigs had telomeric repeats identified on one end (Table S9 in Appendix S1). From these results, we concluded that the four contigs in this genome assembly with telomeres on both ends represented putative chromosomes and the nine contigs with telomeres identified on one end represented chromosome arms. Contig size and gene number did not predict telomere identification, with Contigs 3, 4, 9, and 16 having telomeres identified at both ends, while contigs 1, 5, 6, 8, 10, 12, 14, and 17 had telomeres on one end (Table S9 in Appendix S1). A final summary of all assembly and annotation statistics can be found in Table 1 and Table S10 in Appendix S1.

**Figure 3.**
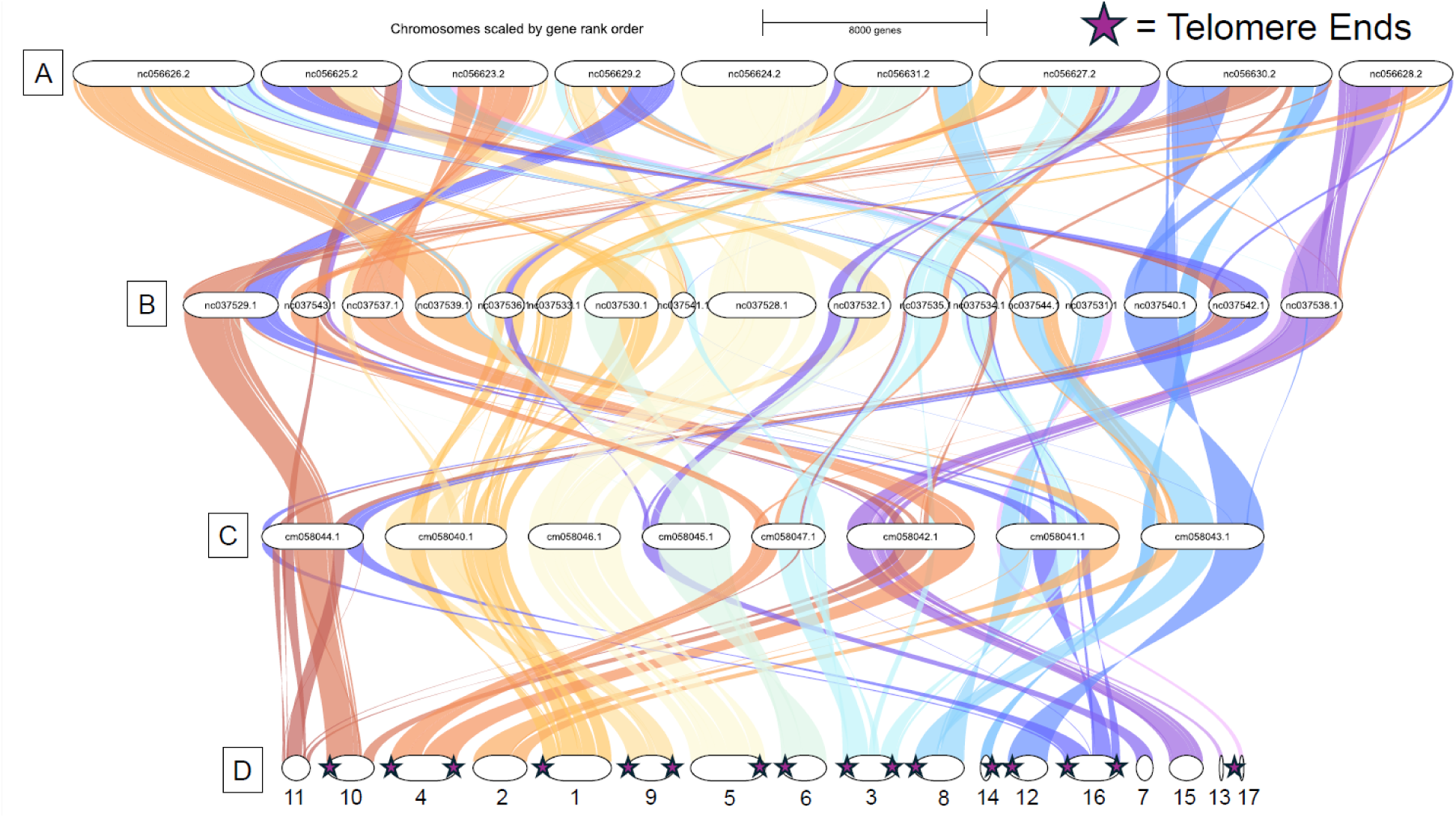
Synteny results from GENESPACE v1.3.1 (Lovell et al. 2022) among assembled genomes of *Centaurea melitensis* and three other Asteraceae: A. *Lactuca sativa* (lettuce), B. *Cynara cardunculus var. scolymus* (artichoke), C. *Centaurea solstitialis* (yellow starthistle), and D. *Centaurea melitensis.* White ovals in (D) represent the 17 largest contigs, each with at least 100 annotated genes. Contigs are numbered based on size (with 1 being the largest and 17 the smallest) and stars represent where telomeric repeats were found using tidk.

**Table 1.** Genome assembly and annotation metrics for *Centaurea melitensis*.

| Datatype | Metric | Value |
| --- | --- | --- |
| Assembly | Total Length | 696,561,998 bp |
|  | # of Contigs | 48 |
|  | Contig N50 | 55,586,964 bp |
|  | Contig N90 | 37,081,860 bp |
|  | Length of Largest Contig | 62,629,239 bp |
|  | # of Putative T2T Chromosomes | 4 |
|  | Genome BUSCO score (eudictos_odb10, n=2326;) | Complete BUSCOs: 98.0% [ single copy: 92.5%, duplicated: 5.5%], fragmented :0.9%, missing: 2.0% |
|  | Inspector QV score | 64.32 |
| Annotation | % of genome occupied by repeats | 65.85% |
|  | # of protein coding genes | 25,157 |
|  | % multi-exonic genes | 72.58% |
|  | Proteome BUSCO score<br>(eudictos_odb10, n=2326;<br>single/duplicated) | 2165 (93.1%) |
|  | OMArk completeness score for<br>asterids clade | 93.57 % completeness |

*Note:* T2T = telomere to telomere; BUSCOs = benchmarking universal single-copy orthologs; QV = quality value; OMArk = orthologous marker assessment

### Synteny

Gene synteny analysis using GENESPACE revealed extensive chromosomal dynamism across *Centaurea melitensis* and three other Asteraceae species: *Lactuca sativa* (lettuce), *Cynara cardunculus var. scolymus* (artichoke), and *Centaurea solstitialis* (yellow starthistle; Figure 3). Across all four species, a total of 24,038 orthogroups were identified. Of these orthogroups, *Centaurea melitensis* shared 17,137 with *L. sativa*, 17,812 with *Cynara cardunculus var. scolymus,* and 18,851 with *Centaurea solstitialis*. However for syntenic pairs, *Centaurea melitensis* shared 14,137 syntenic pairs with its closest relative and *Centaurea solstitialis*, fewer than with the more distantly related *L. sativa* (18,481 pairs) or *Cynara cardunculus var. scolymus* (19,679 pairs). A total of 4,418 shared syntenic blocks and 1,726 syntenic regions were identified across all pairwise comparisons of these genomes.

## DISCUSSION

We successfully assembled and annotated the first reference genome for *Centaurea melitensis* at a near-chromosome level. We report a modest genome size of 795.0 Mbp from flow cytometry, and a 696.6 Mbp genome assembly contained in 48 contigs, including four telomere-to-telomere chromosomes, nine additional chromosome arms terminated by telomeric repeats, as well as a complete chloroplast genome. In comparison to relatives across the Asteraceae with similar chromosome numbers, we find evidence of the evolution of synteny and karyotype, suggesting a dynamic genomic history in the lineage of *C. melitensis*.

Assembly metrics indicated that our assembly is highly complete and largely contiguous. Our >6M reads assembled into just 48 total contigs, including 13 major contigs (>1,000 genes), seven minor gene-containing contigs, and 28 trailing contigs containing no annotated genes. The assembly contained 98% complete eudicot BUSCOs, and more than half of those missing were identified as Pentatricopeptide Repeat (PPR) proteins. PPR proteins are particularly abundant in plants but are commonly missing in assemblies, due to the difficulty of assembling their repetitive tandem 35-amino acid motifs (Tørresen et al. 2019). The *C. melitensis* protein annotations were also largely complete, with 93.1% of the eudicot BUSCO identified.

Our annotations described genome content that is typical among Asteraceae genomes sequenced to date (e.g. Chen et al., 2026; McEvoy et al., 2024; Shen et al., 2018; Xiong et al., 2023). This included 25,157 predicted protein-encoding genes, which is very similar to that reported for *Cynara cardunculus* var. *scolymus* (28,632; Acquadro et al. 2020), another member of the Carduae tribe also used for our synteny comparisons (see below), though somewhat smaller than reported for congener *C. solstitialis* (34,323; Reatini et al., 2024). The total percentage of repetitive elements across all genomic content was 64.2%, which is highly congruent with both *Cynara cardunculus* var. *scolymus* (58.4%; Scaglione et al. 2016) and *C. solstitialis* (63.3%), which are also similar to *C. melitensis* in total genome size. For all three of these genomes, the most common repetitive elements were the Ty1/Copia superfamily (11.52% in *C. melitensis*) and the Gypsy/DIRS1 superfamily (11.08% in *C. melitensis*), a pattern shared across the Asteracaea (e.g. also true of larger and more repeat-rich Asteraceae genomes such as *Lactuca sativa,* Reyes-Chin-Wo et al. 2017, and *Helianthus annuus*; Badouin et al. 2017) though these genomes vary somewhat in which of the two superfamilies is most abundant.

Our reads also contained chloroplast DNA and using these off-target data we were able to assemble a complete chloroplast genome, including the expected large and small single copy regions and two inverted repeats. Chloroplast genome sizes for Asteraceae generally fall between 115,000 bp to 165,000 bp (Daniell et al. 2016), and our *C. melitensis* assembly was within this range at 161,562 bp. It is particularly similar to chloroplast genome assemblies reported for other members of the Cardueae tribe, including *Centaurea cyanus* (152,433 bp, Zhang et al., 2023), and *Carthamnus tinctorius* (153,114 bp; Lu et al. 2016).

Our genome size estimates from flow cytometry, *k*-mer analysis, and the assembly size all suggest a moderately small genome size (under 1 Gbp) relative to the large range for flowering plants (Garcia et al. 2013, Vallès et al. 2013, Vitales et al. 2019). From flow cytometry, we estimated the 1C genome size to be 795.0 Mbp, while our *k*-mer analysis of the PacBio HiFi reads estimated a genome size in the range of 563.81 Mbp (k=17) to 685.17 Mbp (k=121). Our assembly totalled 696.6 Mbp, exceeding the highest *k*-mer estimate. Given that our assembly is not fully gap-less, it is expected that it will be smaller than the flow cytometry estimate, while *k*-mer analyses tend to underestimate genome size relative flow cytometry (Pflug et al. 2020). Flow cytometry estimates themselves also contain various sources of error (Nix et al. 2024), such that our different estimates can all be interpreted as reasonably consistent with a genome size just over 700 Mbp.

Telomere identification in our assembly suggests nine chromosomes in our sequenced individual. Four contigs were bounded by telomeres at both ends and are inferred to be complete chromosomes. An additional nine contigs included a telomere at one end, consistent with at least five additional chromosomes, for a total of nine. Previous reports for *C. melitensis* include nine, 12, and 18 chromosomes (Keil 2012). Nine chromosomes in our sequenced individual would imply that only one telomeric region is missing from the contigs, consistent with an overall highly complete assembly.

We compared gene content and order of our assembly to three other genomes spanning a range of relatedness in the Asteraceae. These included congener *Centaurea solstitialis* (yellow starthistle, 1N = 8; 840 Mbp; Reatini et al. 2024), another member of the Cardueae tribe *Cynara cardunculus* var. *scolymus* (artichoke, 1N = 17, genome size 725 Mbp; Acquadro et al. 2020), and a member of a different tribe (Cichorieae, which shares the subfamily Cichorioideae with the Cardueae) *Lactuca sativa* (lettuce, 1N = 9, genome size 2.5 Gb (Reyes-Chin-Wo et al. 2017). A total of 24,038 orthogroups were identified across these species, a high fraction of total gene content, including 4,418 syntenic blocks and 1,726 syntenic regions. *Centaurea melitensis* shared 17,137 orthogroups with *L. sativa*, 17,812 with C. *cardunculus var. scolymus,* and 18,851 with *C. solstitialis*, consistent with a greater number of shared orthologous genes present and/or detectable as evolutionary distances decreased.

Despite a high fraction of shared gene content, synteny evolution was substantial across these species. Genome structural evolution is well-documented within Asteraceae, even between species which are closely related (Mota et al. 2016, Angulo et al. 2022, Bureš et al. 2023, Feng et al. 2026). Of particular note are large syntenic rearrangements apparent between *C. melitensis* and congener *C. solstitialis* in all four of the putatively complete *C. melitensis* chromosomes bounded by telomeres at both ends. As a result of the substantial syntenic differences between these species, it is not clear from our assembly how the transition between eight and nine chromosomes across these congeners occurred. These patterns suggest rapid and dynamic genome evolution within this clade, with the potential to generate ongoing diversity within *C. melitensis,* as observed in other invasive species systems (Osborne et al. 2026).

This assembly and annotation of the *C. melitensis* genome can be used to facilitate future research and management of this species in multiple ways (North et al., 2021; Hodgins et al., 2025; Kołodziejczyk et al., 2025). The use of reference genomes in combination with population-level genomics has enabled the identification of important demographic processes in invaders, such as changes in effective population size over time (Schoen & Toczydlowski, 2026) and the movement of both beneficial and deleterious alleles across populations in invaded ranges (Putra et al., 2026). For example, the genome assembly of *Amaranthus tuberculatus* Moq. helped determine that resistance to the herbicide glyphosate arose both through the spread of pre-adapted alleles and through the independent evolution of resistant genotypes (Kreiner et al., 2019). Determining whether the success of invasive plants is due to pre-existing standing variation within the native range or is the result of rapid local adaptation in the invaded range is possible by integrating reference genomes with herbarium specimens collected across invaded ranges (Burbano & Gutaker, 2023; Davis, 2023; Kim et al., 2023). Bieker et al. (2022) utilized a genome of *Ambrosia artemisiifolia* L. in combination with genomic resequencing of 308 herbarium specimens to hypothesize that hybridization with closely related species was responsible for its invasion success in Europe. Questions surrounding the evolution of structural variation within the genome and changes in genome size have also been addressed through the use of reference genomes (Yan et al., 2023; Cang et al. 2024). For instance, Wang et al. (2024)’s recent assembly of *Phragmites australis* Cav. uncovered an increased genome size within its invaded range and observed that this genomic expansion was enriched for DNA replication, potentially enabling the fast regeneration and proliferation seen in the invaded range but not in the native range. Finally, determining whether certain advantageous traits are the result of adaptation or plasticity is possible through genomic tools like genome-wide association scans (GWAS) combined with manipulative greenhouse and common garden experiments (Hodgins et al., 2025; North et al., 2021). Recently, Revolinski et al. (2023) utilized a newly created reference genome in combination with GWAS to uncover the pathways responsible for adaptive plasticity within *Bromus tectorum*. Such approaches unlock a detailed understanding of how genetic variants and evolutionary processes contribute to successful invasions, and our reference genome extends these opportunities to *C. melitensis*.

## Acknowledgements

The authors thank the Arizona Genomics Institute for guidance and execution of sequencing and methodology. Support for this research was provided by USDA National Institute of Food and Agriculture awards #2024-67011-43011 to A.J.D., #2024-67012-43394 to J.A.P., and #2023-67013-40169 to K.M.D., and US National Science Foundation awards #1750280 to K.M.D and #2022055 supporting P.C.N.

## Author contributions

All authors were responsible for the conceptual design of this study. All authors were responsible for the writing of this manuscript. A.J.D. led all bioinformatic analyses and was supported by J.A.P, P.C.N, and K.M.D. A.J.D was responsible for all greenhouse work and P.C.N performed the flow cytometry.

## Data availability

The genome assembly for *Centaurea melitensis* has been deposited at NCBI under BioProject PRJNA1433012, BioSample SAMN56353510, and GenBank accession GCA_046736585.1, and will be made public upon acceptance. Raw PacBio HiFi reads, genome assembly, annotation files, and additional results are available on Zenodo (https://zenodo.org/records/18168578). Raw reads will also be deposited in the NCBI Sequence Read Archive (SRA) under the same BioProject following acceptance. A voucher specimen of a sibling of the sequenced individual is deposited at the University of Arizona Herbarium (ARIZ; accession number pending).

## Notes

### Competing Interest Statement

The authors have declared no competing interest.

