## Appendix_Figures_S2 for "Near chromosome-level genome assembly for the invasive annual forb *Centaurea melitensis*"

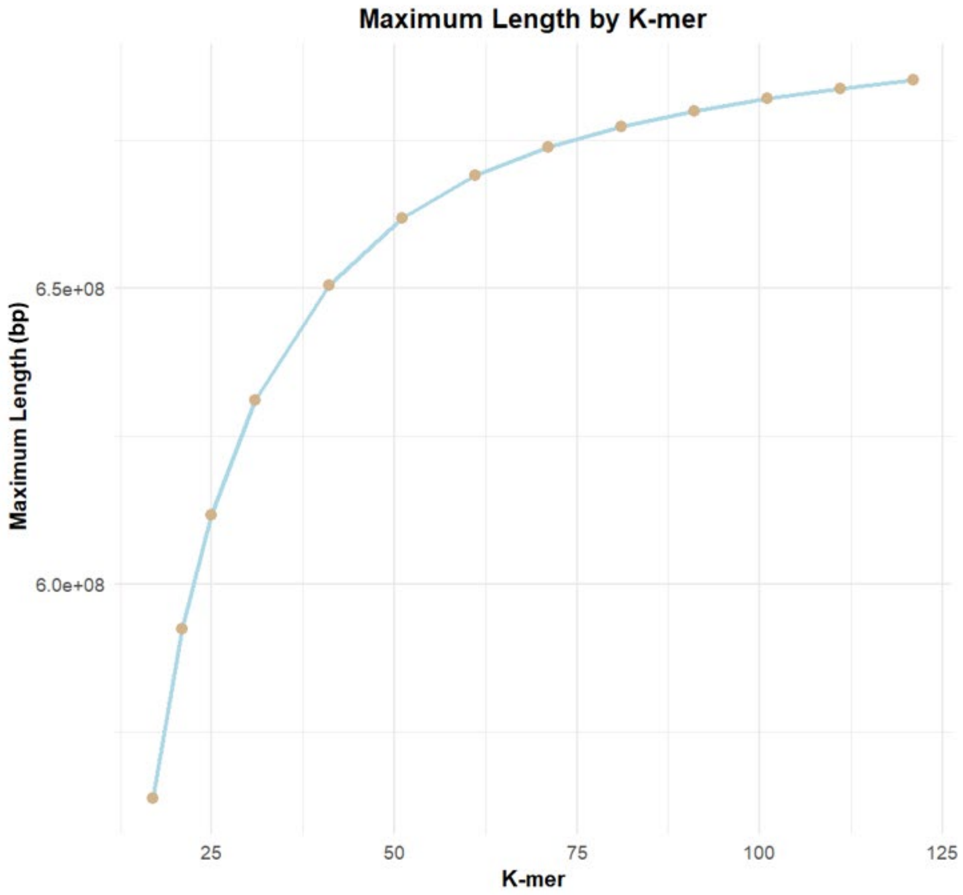


**Figure S1**. Graph of k-mer results using KMC v3.2.4 (Kokot, Dlugosz, and Deorowicz 2017) with default settings to count *k*-mers for *k* = 17, 21, 25, 31, 41, 51, 61, 71, 81, 91, 101, 111, and 121 (shown on the x-axis) and maximum genome length (bp) on the y-axis.

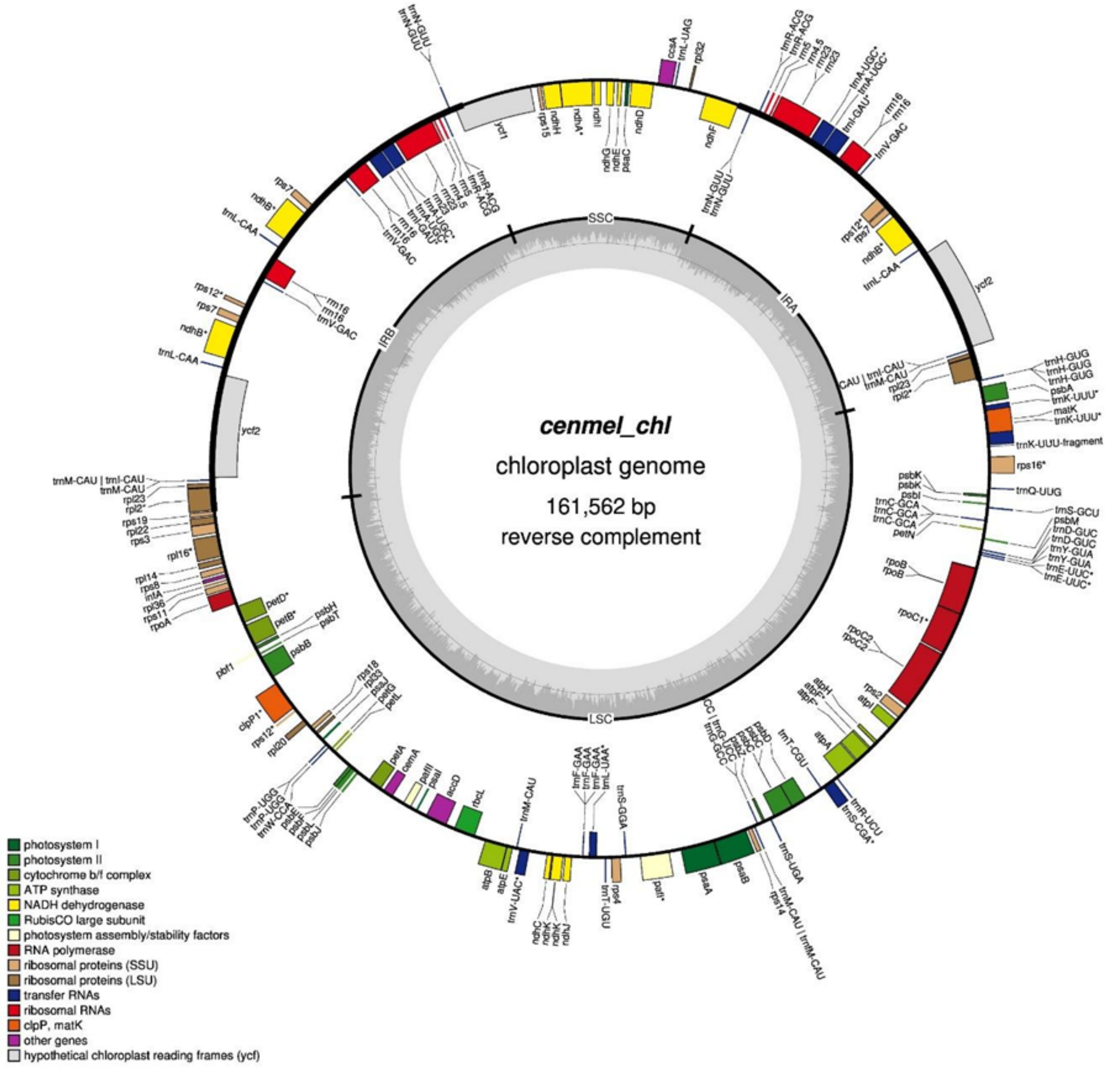


**Figure S2**. Assembled and annotated chloroplast genome of *Centaurea melitensis*.

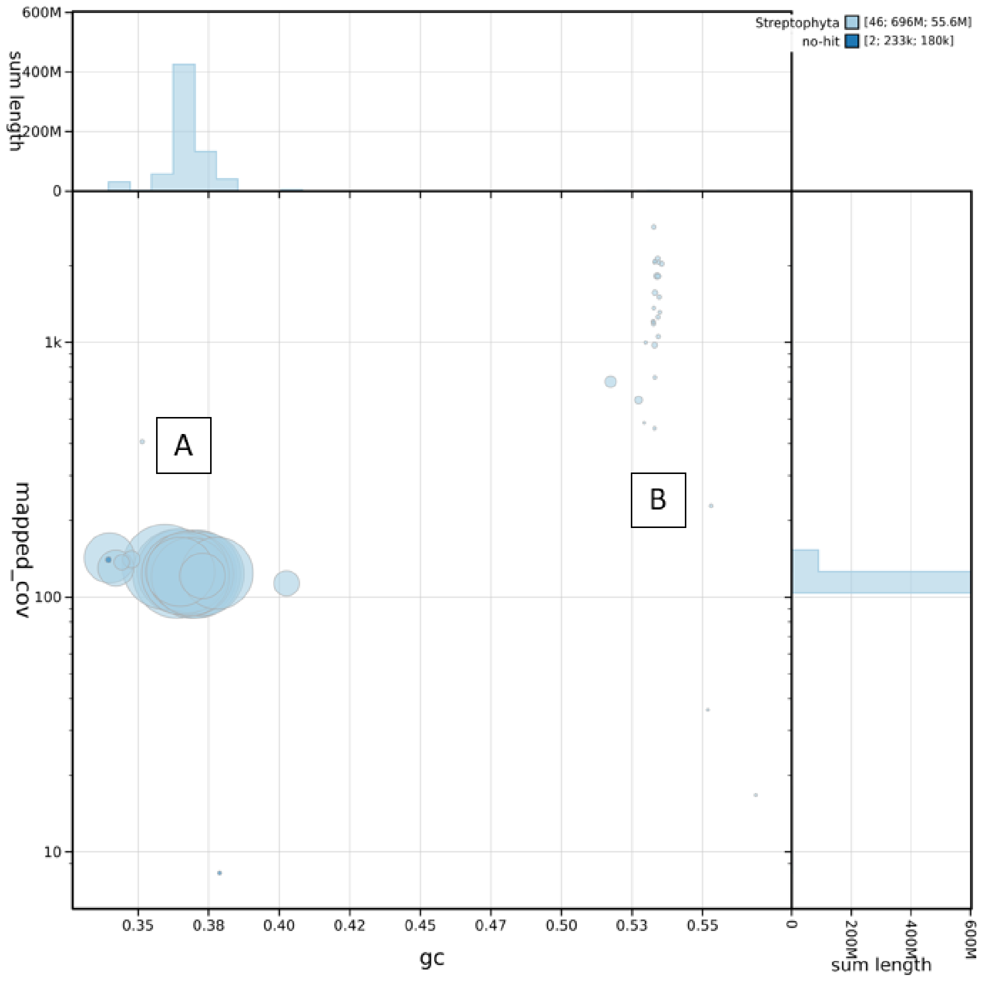


**Figure S3.** Contamination screening using BlobToolKit v4.4.5 (Challis et al., 2020) reveals no contamination in our final *Centaurea melitensis* reference genome assembly. The Blobplot of the assembled genome shows the raw read coverage and GC content for each contig has two distinct regions (labeled A and B), colored by phylum. These correspond to (A) the main contigs of the assembly; (B) contigs containing mitochondrial sequences.
